# Single-axon-resolution intravital imaging reveals a rapid onset form of Wallerian degeneration in the adult neocortex

**DOI:** 10.1101/391425

**Authors:** A.J. Canty, J.S. Jackson, L. Huang, A. Trabalza, C. Bass, G. Little, V De Paola

## Abstract

Despite the widespread occurrence of axon degeneration in the injured and diseased nervous system, the mechanisms of the degenerative process remain incompletely understood. In particular, the factors that regulate how individual axons degenerate within their native environment in the mammalian brain are unknown. Longitudinal imaging of >120 individually injured cortical axons revealed a threshold length below which injured axons undergo a rapid-onset form of Wallerian degeneration (ROWD). ROWD consistently starts 10 times earlier and is executed 4 times slower than classic Wallerian degeneration (WD). ROWD is dependent on synaptic density, unlike WD, but is independent of axon complexity. Finally, we provide both pharmacological and genetic evidence that a Nicotinamide Adenine Dinucleotide (NAD^+^)-dependent pathway controls cortical axon ROWD independent of transcription in the damaged neurons. Thus, our data redefine the therapeutic window for intervention to maintain neurological function in injured cortical neurons, and support the use of *in vivo* optical imaging to gain unique insights into the mechanisms of axon degeneration in the brain.

## INTRODUCTION

Axon degeneration leads to loss of connectivity and brain functions in neurodevelopmental diseases, trauma including head injury and stroke, and age-related conditions such as in the early stages of Alzheimer’s disease. Understanding the mechanisms of axon degeneration to either prevent or delay it, therefore, represents a promising avenue to develop effective therapies for these brain disorders (*1*). Existing, widely used experimental models, however, are focussed on injured myelinated fibres of the optic nerve (*2*), spinal cord (*3*) and peripheral nervous system (*4*), which are not the main axon types affected in brain diseases. Therefore, whether similar mechanisms underlie axon degeneration in the adult brain grey matter is unknown. In particular, we lack critical information on unmyelinated cortical axons, a fibre system vulnerable in many of the afore-mentioned disorders.

Axonal injury is followed by degeneration of the distal part of the axon that is detached from the soma. This tightly controlled process of axon self-elimination following injury is termed WD (*5*), and has been intensively studied and characterised with the aim of devising new strategies to treat neurodegenerative and other diseases (*1*), which typically manifest with prominent local axon degeneration preceding cell body loss. Different molecular mechanisms control axonal and cell body degeneration (e.g. apoptosis) (*6, 7*). In mouse models of neurodegeneration, preventing apoptotic cell death has little or no influence on the axonal degeneration kinetics and disease duration (*8–12*). Preventing or delaying axon degeneration could not only slow down the course of neurodegenerative conditions (*13–15*), but also delay the onset of loss of functionality by maintaining or stabilising the integrity of the neurons (*16*), which are postsynaptic to the damaged axons.

There are similarities between the morphological and functional features of WD and the axon pathology typical of neurodegenerative disease, e.g. common cellular pathways (*17*) (such as autophagy or altered axonal transport (*18*)), and stages (such as axon fragmentation), but also differences, including distinct molecular pathways (*19*). After physical injury, the portion of the axon disconnected from the soma (also called the distal axon) undergoes calcium influx-mediated morphological changes (*20*), including thinning, beading and resealing of the cut end, and is initially able to transmit action potentials (*21, 22*). Calcium overload drives mitochondria-mediated reactive oxygen species damage (*23*) and activates the protease calpain, which degrades neurofilaments, so that after 24-48 hours (*24–27*) the distal segment begins to fragment and is eventually removed by resident glial cells (*5, 28, 29*). The reason for such a protracted and variable lag phase before fragmentation has remained unclear, although its duration is regulated by the ubiquitination machinery in zebrafish larvae (*30*). Previous studies have suggested that the progression of axon degeneration depends on the length of the disconnected segment, with longer segments taking longer to degenerate than shorter ones (*20, 27*). This length-based mechanism is consistent with the presence of axon protective factors such as NAD^+^ or the NAD^+^ synthesising enzymes, which when depleted from the axon trigger the process of fragmentation (*31–34*), with longer disconnected segments having inherently higher levels of such protective factors compared to shorter ones. This idea has been difficult to test because of the challenge of measuring the length of transected axons before the lesion in the living brain.

After injury, the axonal membrane is more resilient to degeneration compared to cytosolic and cytoskeletal axonal components, which are affected first (*17, 32*). *In vivo* imaging studies in the murine spinal cord (*35*) and optic nerve (*2, 36*) have also described the process of acute axonal degeneration (AAD), occurring within minutes of injury. AAD affects 200-300 μm on both severed endings of myelinated axons (i.e. both the proximal and distal sides), followed by the removal of the rest of the distal portion after 24-48 hours by WD (*35*). Whether cortical axons also undergo AAD-like processes still remains to be established. In addition, because these and most previous studies used cytosolic or cytoskeletal reporters (*2, 30, 35, 37–42*), the kinetics and mechanisms of axolemmal fragmentation remain unexplored.

Axons constitute the largest neuronal compartment, surpassing the size of the cell body 1000-fold, in both length and surface area (*43*). They can extend for several tens of cm in the mouse brain (*44*) and branch thousands of times to reach all their targets (*45*). As a result, it has not been possible to follow the degeneration process for individual unmyelinated axonal arbours in the highly interconnected cortical grey matter, leaving the precise spatiotemporal progression and synaptic mechanisms (*46*) uncharted.

Here, we use *in vivo* multiphoton microscopy to capture for the first time the cellular dynamics of cortical axon degeneration for up to 3 days post injury. We report four main findings: First, we identify a threshold length (~ 600 μm) below which damaged axonal segments consistently and selectively undergo ROWD, which starts 10 times earlier and is executed 4 times slower than WD. Second, ROWD depends on synaptic density but not on the complexity of the axonal arbour. Third, AAD is negligible on cortical unmyelinated axons. Finally, a local NAD^+^-dependent pathway regulates ROWD, independently of gene transcription in the damaged neuron.

## RESULTS

### Single-axon-resolution imaging of Wallerian degeneration in the living brain

In order to study the progression of axonal injury at single axon resolution over several days in the otherwise intact brain, we implanted cranial windows over the somatosensory cortex and used a well-established microlesion paradigm based on femtosecond laser-mediated subcellular ablation (*38, 47–53*). High-energy lasers can be non-invasively directed with high precision to specific axonal segments, causing localised axonal breakage (*54*).

To monitor the integrity of the axonal membrane in real-time (*17, 20*), we opted for a membrane-targeted form of GFP under the Thy1 promoter (*55*), which drives expression of transgenes predominantly in excitatory cells of cortical layers 2/3/5/6 and thalamic origin.

### A Rapid Onset form of Wallerian Degeneration

Individual cortical axons of known length from the ending (range 0.12 – 1.5 mm *in vivo* and 0.12 – 5 mm *in vitro*) were identified and transected (See methods, **Fig. 1**) as previously shown (*48–50*).

**Figure 1.**
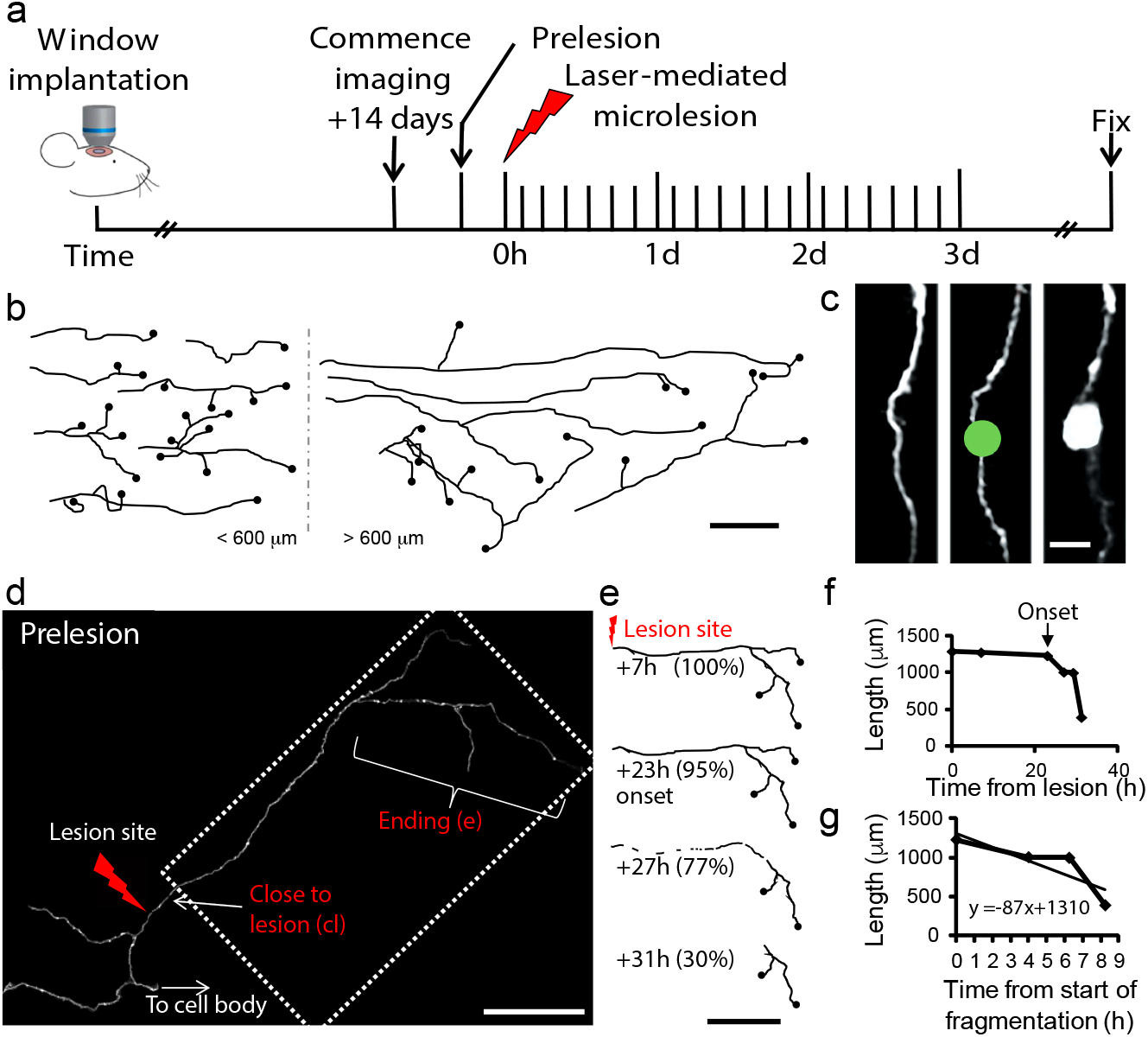
Single-axon-resolution imaging of Wallerian degeneration in the living brain. **(a)** Timeline of the experiments to track non-myelinated axon degeneration *in vivo*. **(b)** Representative drawings of axon segments before lesioning used throughout the study. Black filled circles indicate axon endings. Scale bar: 150 μm. **(c)** Higher magnification image of a small region of axon ready for lesioning (left). A binary mask (4 μm) is placed over the axon (middle) and a focused laser beam used to induce axotomy (right). Proximal is up and distal is down. Scale bar: 5 μm. **(d)** A representative axon before lesioning (total length 1289 μm). The portion in the dotted box is shown as a drawing in (e). Scale bar: 200 μm. **(e)** Representative drawings of the disconnected portion of the axon in the dotted box in (d). *In vivo* two-photon imaging revealed the process of axonal degeneration in the intact brain over selected time points. Lightning bolt is lesion site. Scale bar: 300 μm. **(f)** Representative example of quantification of the degeneration onset for the axon in d-e (23 hours). **(g)** Representative example of quantification of the fragmentation rate for the axon in d-e (1.45 μm/min).

Surprisingly, we found that disconnected axons < ~ 600 μm started the degeneration process by fragmenting much earlier than longer segments (> ~ 600 μm) (**Fig. 2a, b**). Indeed, relating the length and onset times for 30 transected cortical axons suggested the presence of two types of responses (**Fig. 2c**).

**Figure 2.**
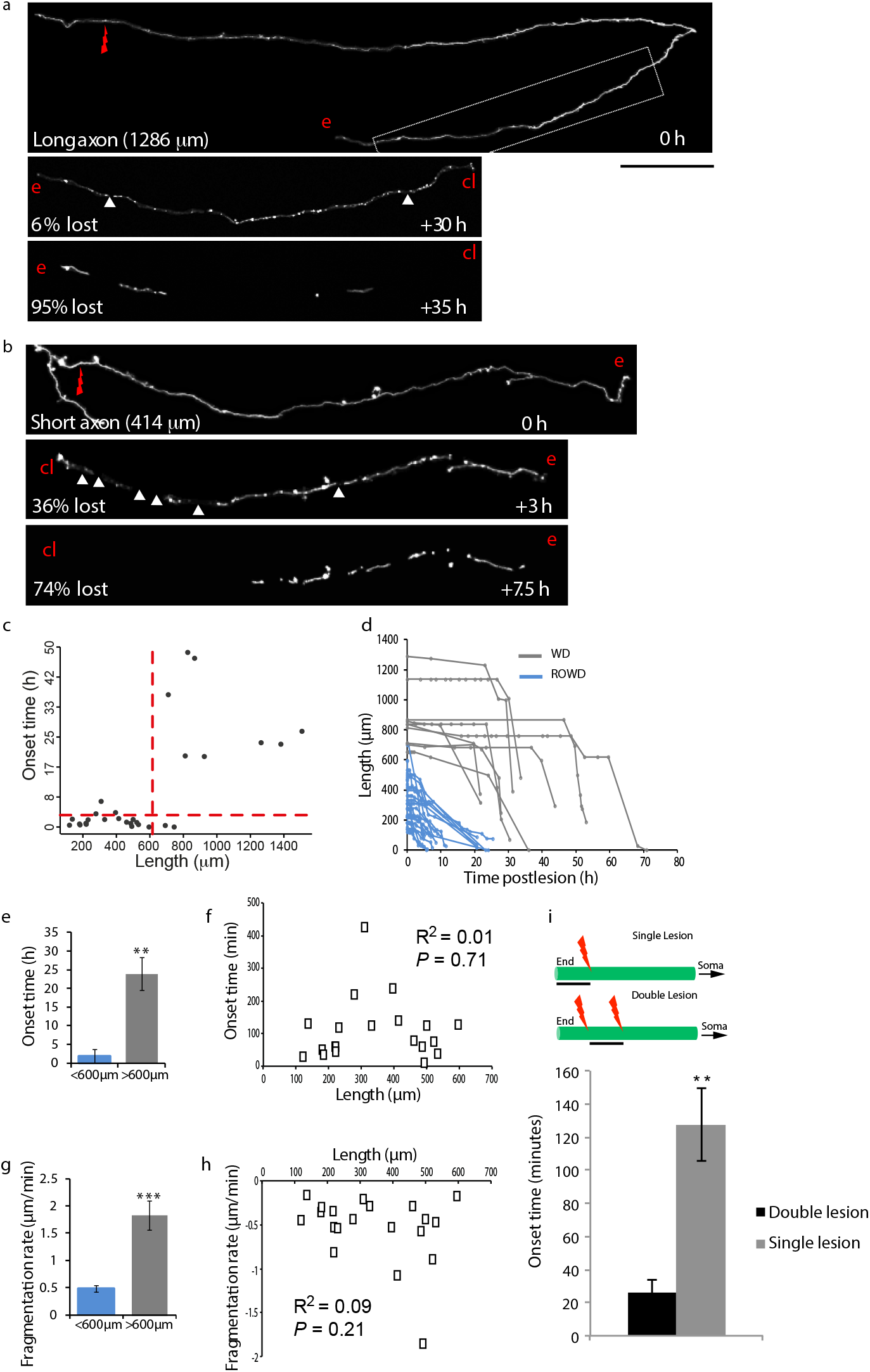
A rapid onset form of Wallerian degeneration in the adult cortex. (**a**) Representative images of an axon undergoing WD. Scale bar: 115 μm in the 0 h panel and 80 μm in the two panels below (30 and 35 h). (**b**) Representative images of the terminal portion of an axon undergoing ROWD. Scale bar: 70 μm. (**c**) Scatter plot indicating the presence of two populations with different fragmentation onsets. Y-axis red line = 3h; x-axis red line = 615 μm (see text for details). (**d**) Axon length imaged *in vivo* (n = 30) over the time points indicated revealing an early-onset form of axon degeneration (blue lines). (**e**) Earlier onset time of ROWD (blue bar) compared to WD (grey bar). (**f**) No correlation between ROWD onset time and length (n = 20). (**g**) Slower fragmentation rate of ROWD (blue bar) compared to WD (grey bar). (**h**) No correlation between ROWD fragmentation rate and length (n = 20). (**i**) The average onset for single lesions is significantly higher than for double lesions, comparing cut segments of equal length (scale bars). This experiment supports a protective factor level cut-off based model of cortical axon degeneration (see text for details). Spearman’s correlation analysis and Mann Whitney U test. Red arrows mark lesion site. cl = close to lesion site end, e = end of severed axon. Mean ± SEM. Mann Whitney U test. ** *P* < 0.01, *** *P* < 0.001.

Following this observation of bimodality within the length and onset time data (**Fig. 2c**) we sought to rigorously test the existence of two populations. The two Gaussian components of these bimodal distributions were fitted using the R mixtools packages (Version 1.0.2). The mean and standard deviations were calculated for these modelled Gaussian components, and the upper 95% confidence intervals of the components with the lower mean within both the onset time and length measurements identified. This allowed for the classification of a subset of data points as significant outliers (*P* < 0.05, bimodality test) to the lower components of both onset time (i.e. points above 3.3 h, red dashed line y-axis intercept, **Fig. 2c**) and length (points above 615 μm, red dashed line x-axis intercept, **Fig. 2c**). This subclass of data points was found to have significantly higher mean onset times compared to the rest of the data (P < 0.01, Wilcoxon rank sum test). Indeed, tracking individual injured axons continuously up to 70 hours (~ 3 days) at < 4h intervals (see Methods) to capture the onset of fragmentation, showed that longer transected axonal segments (typically > 600 μm from the end; mean = 973 μm, *n* = 10), despite being lower in numbers, consistently initiated fragmentation after 24-48 hours in line with the slow onset described in the WD literature (8 out of 10 examples, **Fig. 2a, d, e**). In contrast, shorter transected axonal segments (typically < 600 μm from the end; mean = 341 μm, *n* = 20) consistently initiated fragmentation on average 10 times earlier than the longer portions (19 out of 20 examples; **Fig. 2b, d, e**, < 600 μm, onset time = 1.7 ± 0.4 hours, *n* = 20; > 600 μm, onset time = 24.5 ± 4.10 hours, *n* = 10, *P* < 0.01), via a process we termed ROWD.

However, despite falling into two clear groups (short-endings < 600 μm undergoing ROWD or long-endings > 600 μm undergoing WD), we found no correlation between the time of onset of fragmentation and the actual length of the disconnected axon endings within each group (**Fig. 2f**, < 600 μm undergoing ROWD onset time versus length, R^2^ = 0.01; *P* = 0.71; **Supplementary Fig. 1**, > 600 μm undergoing WD, *P* > 0.05), suggesting the presence of a cut-off-dependent mechanism. These results are in contrast to the previously postulated linear length-based model to explain the onset of degeneration (*20, 27, 30*), according to which the longer the disconnected segment, the longer the degeneration onset time, but consistent with two alternative cut-off based models. According to the first, for axotomised segments < ~ 600 μm, the amount of a protective factor (e.g. NAD, a diffusible molecule, or Nmnat2, one of the neuroprotective NAD synthesising enzymes (*31, 56, 57*)) would decrease below a critical threshold level before the broken membrane can reseal, triggering ROWD. For segments > 600 μm, the resealing of the membrane from the cut end would happen before the depletion of the protective factor is able to trigger ROWD. According to the second model, a critical threshold level of a WD trigger such as calcium influx would be rapidly reached in segments < ~ 600 μm, because of their limited calcium buffer capacity. The first model (protective factor level cut-off model) predicts that axonal segments with two cut ends will have earlier onset time than equal length segments having only one cut end, as the protective factor would have two depletion routes, instead of one. According to the second model (WD trigger amount cut-off model), however, there should not be any difference between the degeneration time of segments of equal length, irrespective of whether they have one or two cut ends. This is because the calcium increase in the axoplasm, i.e. the putative WD trigger, is mediated by calcium channels and/or internal organelles distributed along the length of the axon (*58, 59*), and therefore expected to be the same for segments of equal length. To distinguish between the two models, we performed experiments in which two lesions were made along the axon. Segments < 600 μm with two cut ends had an earlier degeneration onset than equal length segments with only one cut end, supporting the protective factor cut-off based model (**Fig. 2i**, single lesion onset = 127 ± 22 minutes; double lesion onset = 26 ± 8 minutes; *P* = 0.001; single lesion segment length = 179 ± 9 μm, *n* = 8 axons; double lesion segment length = 200 ± 16 μm, *n* = 10 axons; *P* = 0.2).

To further elucidate the mechanisms of ROWD and WD, we compared the fragmentation rates once onset was detected. Surprisingly, the rate of fragmentation for shorter transected axonal segments (< 600 μm from the end) was significantly slower than for longer transected axonal segments (**Fig. 2g**, < 600 μm, 0.5 ± 0.1 μm/min, *n* = 20; > 600 μm, 1.9 ± 0.2 μm/min, *n* = 10, *P* < 0.01). Within each group (short: < 600 μm or long endings: > 600 μm) there was no correlation between the length of the disconnected axon segments and the fragmentation rate (**Fig. 2h**, < 600 μm fragmentation rate versus length, R^2^ = 0.09, *P* = 0.21; **Supplementary Fig. 1**, > 600 μm, *P* > 0.05), again ruling out a linear length-based mechanism to explain the degeneration kinetics, supporting instead a cutoff-based mechanism.

### AAD is negligible in injured cortical axons *in vivo*

More frequent regular imaging over time revealed little evidence of cortical axon fragmentation occurring within the first 60 minutes post-lesion, either proximal (*50*) or distal to the lesion site (**Supplementary Fig. 2**, **Supplementary Movie 1**), with all but one disconnected axonal segments exhibiting a longer delay in the onset of fragmentation (**Supplementary Fig. 2a and c**; 25 out of 26, 96%) (**Fig. 2, Table 1**).

**Table 1.** Axon degeneration modalities.

| **Degeneration mechanism** | **Marker** | **Fragmentation onset** | **Fragmentation rate** | **Axon length affected** |
| --- | --- | --- | --- | --- |
| Acute Axonal Degeneration | Cytosolic GFP | 2-60 min | 40-60 $\mu\text{m}/\text{min}$ | 200-300 $\mu\text{m}$, both distally and proximally (2, 35) (this study) |
| Rapid Onset Wallerian Degeneration (< 600 $\mu\text{m}$ cortical axons) | Membrane GFP | $1.7 \pm 0.4 \text{ h}$ | $0.5 \pm 0.1 \mu\text{m}/\text{min}^*$ | < 600 $\mu\text{m}$, only distally (this study) |
| Wallerian Degeneration (cortical axons) | Membrane GFP | 24.5 ± 4.1 h | 1.9 ± 0.2 $\mu\text{m}/\text{min}^*$ | > 600 $\mu\text{m}$, only distally (this study) |
| Wallerian Degeneration (spinal cord) | Cytosolic GFP | ~24-30 h | 7-366 $\mu\text{m}/\text{min}$ | Several mm (62);(35, 63), only distally |

\*See methods for differences between our sampling protocol and previous literature.

The absence of acute membrane fragmentation in the unmyelinated cortical grey matter is in contrast with the process of AAD as described in the myelinated fibres of spinal cord (*35*) and optic nerve (*2*), where axonal fragmentation occurred on both sides of the lesion within few minutes after the injury. This difference could be due to different lesion methods (laser versus mechanical axotomy), markers used (axonal membrane versus cytosolic) or intrinsic characteristics of the different neuronal populations in the nervous system (cortex versus spinal cord/optic nerve). To investigate these possibilities we performed laser axotomies in both the ascending dorsal white matter tracts in the murine spinal cord (where AAD was first described (*35*)) and in cortical axons using either cytosolic (GFP-M line (*60*)) or axolemmal GFP markers (L15 line (*55, 61*)), expressed under the Thy 1.2 promoter. *In vivo* time-lapse imaging of individual axons showed that while the axoplasm fragmented within minutes and in proximity of the spinal cord lesion (axoplasm degeneration onset = 29.6 ± 4 min; range 16 - 56 min, *n* = 9 axons), consistent with the previously described onset of AAD (*38*), the membrane near the lesion site remained intact for up to 4 hours (axolemmal onset = 139.9 ± 22 min; range 58 - 228 min, *n* = 9 axons; **Supplementary Fig. 3**; *P* < 0.001 compared to axoplasmic onset). In addition, in cortical axons the axoplasm of the distal disconnected portion did not fragment but appeared retracting from the lesion site over the first 60 min postlesion (**Supplementary Fig. 2b-c**; mean distance from lesion site = 11.1 ± 1.9 μm; range 3 - 21 μm; *n* = 10 axons), likely due to leakage of GFP from the cut end immediately after the lesion, revealing a surprising difference between spinal cord and cortical axons in their initial response to injury.

Unlike a previous study in zebrafish larvae (*30*), measuring cortical axon fragmentation with cytosolic markers in our study yielded significantly different rates than with membrane markers (3.4 ± 0.7 μm / min versus 0.5 ± 0.1 μm / min, respectively; *P* < 0.05). These fragmentation rates are also much slower than the ones measured from previous studies (*27, 35*), a finding explained by the longer sampling intervals used in our chronic imaging protocols (see methods for details on the fragmentation rate measurements). These results provide *in vivo* evidence that cytosolic and membrane markers report different aspects of axonal degeneration, and that AAD in the spinal cord (*35*) defines a process of axoplasmic fragmentation uncoupled from axolemmal breakage.

Taken together, we describe novel dynamics of cortical axon degeneration following physical injury *in vivo*. We identify a threshold length (~ 600 μm) below which the lag phase before cortical axon fragmentation is one order of magnitude shorter and the fragmentation rate significantly slower than WD. In addition, compared to AAD (*2, 35*), ROWD removes the entire severed distal axon rather than just a portion of it adjacent to the lesion site, and does not occur on the proximal axonal segment (i.e. the portion still connected to the cell body), revealing spatiotemporal differences between ROWD and WD/AAD (**Table 1**).

### ROWD depends on synaptic density but not axonal arbour complexity

We then asked how the density of synapses along cortical axons and the complexity of the arborisation affect ROWD and WD. We reasoned that axons with a relatively high density of synaptic contacts may take longer to degenerate than axons with lower synaptic density due to the additional time required to disassemble the synaptic machinery (*32*). The synaptic density (range: 0.0048-0.0939 synapses/μm) of 30 axons was quantified on the distal portion prior to the laser micro-lesion and correlated with the onset and rate of fragmentation of the same segments (**Fig. 3**). Surprisingly, while for segments > 600 μm undergoing WD, synaptic density did not significantly correlate with either the onset (P > 0.05; **Fig. 3d**) or rate of fragmentation (P > 0.05; **Fig. 3f**), for segments < 600 μm undergoing ROWD, synaptic density significantly correlated with the onset (R^2^ = 0.30, *P* = 0.01; **Fig. 3c**), but not with the rate of fragmentation (R^2^ = 0.03, *P* = 0.49; **Fig. 3e**).

**Figure 3.**
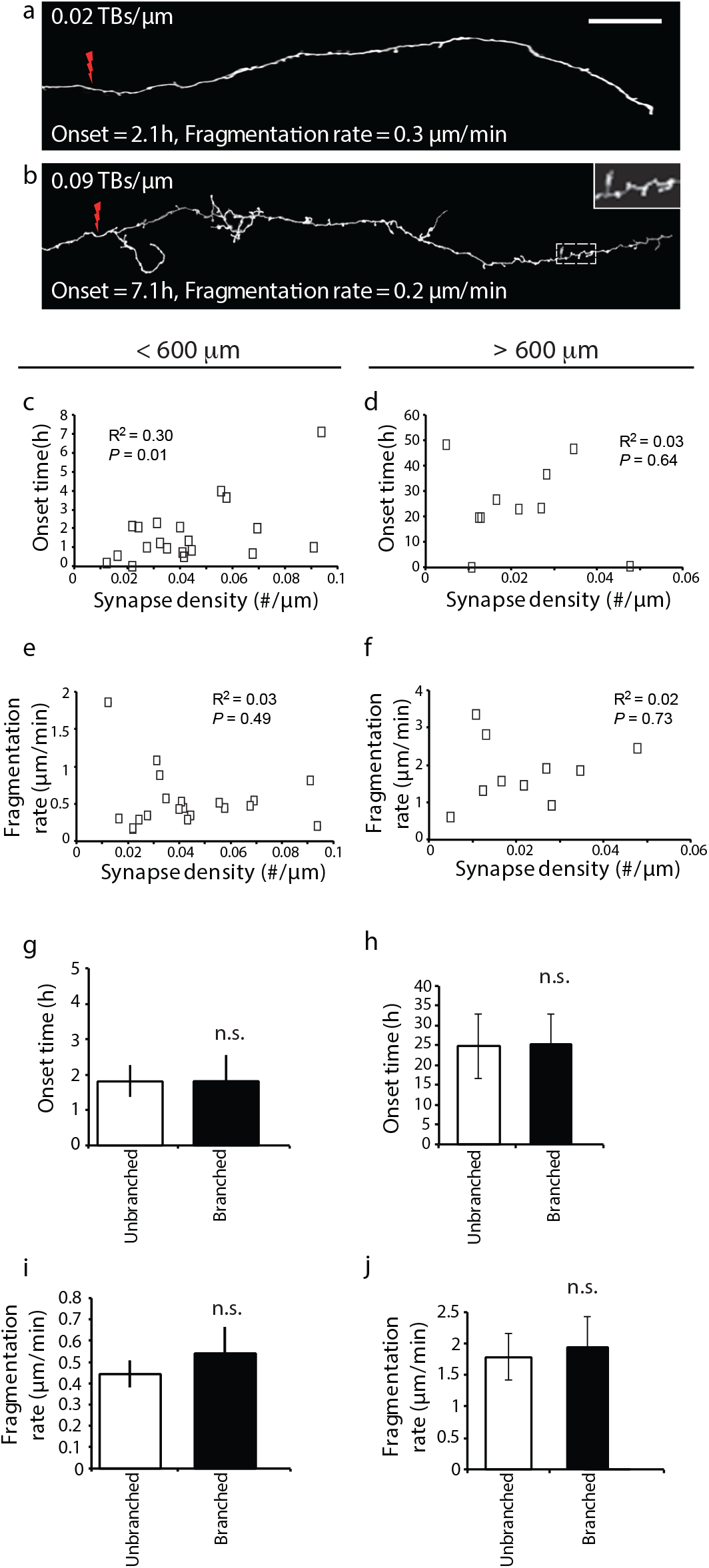
ROWD depends on synaptic density but not axonal arbour complexity. (**a**) Representative image of an axon with relatively low synaptic density. Lightning bolt marks the lesion site. Scale bar: 40 μm. (**b**) Representative image of an axon with relatively high synaptic density. Segments < 600 μm (c, e, g, i). Segments > 600 μm (d, f, h, j). (**c**) Correlation between onset time and synaptic density for segments undergoing ROWD. (**d**) No correlation between onset time and synaptic density for segments undergoing WD. (**e**) No correlation between fragmentation rate and synaptic density for segments undergoing ROWD. (**f**) No correlation between fragmentation rate and synaptic density for segments undergoing WD. (**g, h**) Fragmentation onset times for unbranched and branched axons. (**i, j**) Fragmentation rates for unbranched and branched axons. Mean ± SEM. Spearman’s correlation analysis and Mann Whitney U test values considered significant at *P* < 0.05.

These data suggest that, as initially hypothesised, for axons undergoing ROWD the depletion rate of the protective factor may be dependent on the density of synaptic components, so that the higher the density of synaptic contacts along the axon, the longer the lag phase before fragmentation. Once the execution phase starts, however, the density of synaptic elements does not affect the kinetics of fragmentation. Moreover, branched and unbranched axons had similar onset times (**Fig. 3g**, h; unbranched < 600 μm, 1.80 ± 0.45 h, *n* = 15; branched < 600 μm, 1.81 ± 0.74h, *n* = 5, *P* > 0.05; unbranched > 600 μm, 24.7 ± 8.0h, *n* = 5; branched > 600 μm, 24.3 ± 7.7h, *n* = 5, *P* > 0.05) and fragmentation rates (**Fig. 3i**, j; unbranched < 600 μm, 0.44 ± 0.06 min; branched < 600 μm, 0.54 ± 0.12 min, *P* > 0.05; unbranched > 600 μm, 1.81 ± 0.39 min, *n* = 5; branched > 600 μm, 1.92 ± 0.49 min, *n* = 5, *P* > 0.05), suggesting that axon complexity does not play a major role in the degeneration process.

### ROWD is controlled by a local NAD^+^ dependent pathway *in vitro* and *in vivo*

To uncover the mechanism of cortical axon ROWD we assessed NAD^+^-dependent pathways, already implicated in WD and AAD (*35, 64, 65*) and in modulating axon degeneration in several animal models of disease (*1*). We used the Wallerian degeneration slow (Wld^S^) mouse line overexpressing a stable version of the NAD^+^ synthesizing enzyme NMNAT1, which prevents the NAD^+^ level decrease after axotomy (*65, 66*). Animals with the Wld^S^ mutation have delayed onset of WD in both the peripheral and central nervous system (*67, 68*). Wld^S^ animals were crossed with the Thy1-L15 (**WT**) line to produce heterozygous Wld^S^-Thy1-L15 progeny. Severed axons < 600 μm from the end in both Wld^S^-Thy1-L15 (n = 24) and WT animals (n = 20) were tracked (**Fig. 4a-d**). While there was no significant difference in fragmentation rate (**Fig. 4d**; WT, 0.5 ± 0.1 μm/min; Wld^S^-Thy1-L15, 1.7 ± 0.54 μm/min, *P* > 0.05), Wld^S^-Thy-L15 animals showed a delayed onset of fragmentation (**Fig. 4c**), with an overall significant increase in the onset time compared to WT mice (WT, 1.7 ± 0.4 hours; Wld^S^-Thy1-L15, 7.6 ± 3.7 hours, *P* < 0.01), indicating that similarly to WD and AAD, also ROWD is regulated by NAD-dependent pathways.

**Figure 4.**
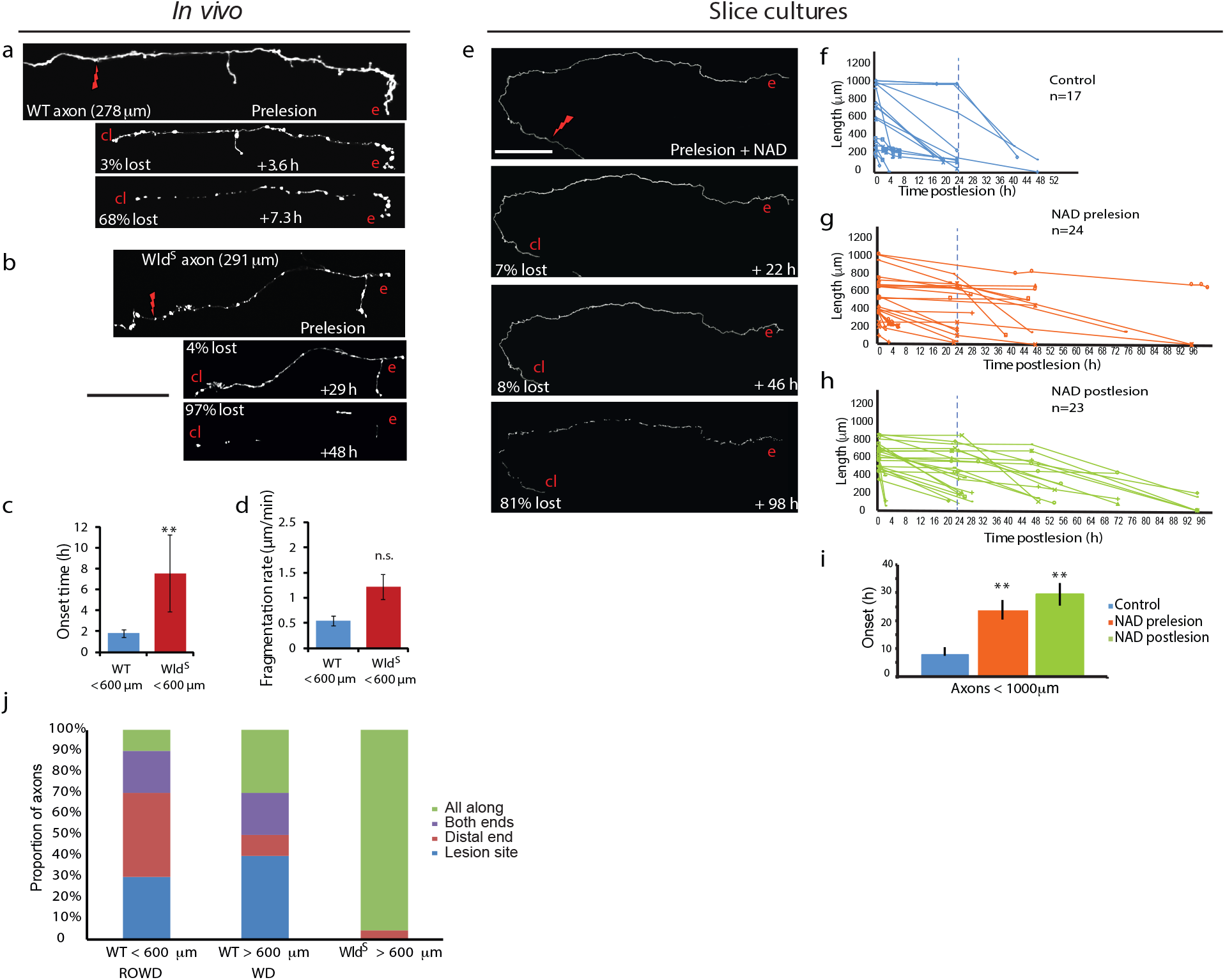
ROWD is controlled by a local NAD^+^ dependent pathway *in vitro* and *in vivo*. (**a-d**) Wld^S^ delays ROWD *in vivo*. (**a**) Representative example of ROWD in WT animals imaged *in vivo*. (**b**) Representative images of a Wld^S^–Thy1-L15 axon where a comparable distal length had been cut off *in vivo*. Scale bar in A and B: 70 μm. (**c**) Wld^S^ group had significantly longer onset times than WT. (**d**) The fragmentation rates were not significantly different in the WT and Wld^S^ groups. (**e-i**) NAD^+^ delays ROWD *in vitro*. (**e**) Representative images of the degeneration of an axon in a slice culture treated with NAD^+^. Scale bar: 100 μm. (**f-i**) Degeneration of *in vitro* lesioned axons with no treatment (control, **f**), with addition of NAD^+^ prelesion (**g**) and postlesion (**h**). Blue dotted line shows the 24h time point. (**i**) Axons are protected by the addition of NAD^+^ pre- and postlesion treatment. Red lightning bolts mark lesion site. cl = close to lesion end, e = distal end of severed axon. (**j**) Quantification of spatial location of first axon fragments. Mean ± SEM. Mann Whitney U test. ** *P* < 0.01.

To learn more about the NAD^+^-dependent regulation we used cortical explant cultures, amenable to pharmacological manipulation (*69*). Axotomized terminal axons labeled with membrane-bound GFP (mean length = 613 ± 80 μm, *n* = 17 axons from 6 brains) undertook ROWD with comparable fragmentation onset according to the dynamics described in *in vivo* experiments (**Fig. 4e-f**; onset = 1.78 ± 0.18 h, 9 axons < 600 μm; onset = 17.6 ± 0.18 h, 8 axons 600-1000 μm; *P* > 0.05 compared to *in vivo* measurements). Fragmentation rates were not calculated for the *in vitro* experiments as the room temperature imaging conditions confounded the fragmentation kinetics (*62*). The protective effect of NAD may be due to a local action on the cut distal axon or to its role in modulating gene expression in the nucleus (*64*).

To distinguish between these possibilities, we first applied 1 mM NAD^+^ 24 hours before the lesions, which resulted in robust delay in ROWD onset (**Fig. 4g**, i; mean length = 521 ± 51 μm, onset = 23.9 ± 3.5h, *n* = 24; *P* < 0.01 compared to untreated). To rule out any transcription-dependent NAD^+^ effects in the damaged neurons (*64*), we also applied NAD^+^ immediately after the axonal cut. In this case, NAD^+^ was also able to delay the onset of degeneration when applied after the lesion (**Fig. 4h**, i; length = 613 μm, onset = 30.4 ± 4.2h, *n* = 23; *P* < 0.01). These data suggest that the NAD^+^ action is local, in agreement with earlier work (*65, 70, 71*), rather than requiring gene expression in the axotomised neurons as previously shown for WD in dissociated neuronal cultures (*64*).

Finally, we determined where the fragmentation first began along injured cortical axons. Out of 30 axons, 10 started fragmenting from the lesion site, 9 from the end furthest from the lesion, 5 along the entire length at the same time, 3 from the middle and 3 from both ends simultaneously, with different regions displaying comparable onset (close to lesion site, onset time = 7.3 ± 2.4 hours, *n* = 26; distal end, onset time = 10.4 ± 2.9 hours, *n* = 26, *P* > 0.05) and fragmentation speed (close to lesion end, 0.19 ± 0.035 μm/min, *n* = 16; distal end, 0.19 ± 0.038 μm/min, *n* = 16, *P* > 0.05). Intriguingly, in unmyelinated cortical axons, ROWD is significantly more likely to start at the distal end than WD (**Fig. 4j**; < 600 μm ROWD axons, 40%, 30%, 20% and 10% started fragmenting at the distal end, close to the lesion end, both ends simultaneously or all along the axon, respectively; > 600 μm WD axons, 10%, 40%, 20% and 30% started fragmenting at the distal end, close to the lesion end, both ends simultaneously or all along the axon, respectively; Chi squared test, *P* < 0.001), highlighting further differences between ROWD and WD.

## DISCUSSION

What are the mechanisms of axon degeneration in the adult injured brain? In this study we used *in vivo* imaging through a cranial window and laser-mediated lesions to track the full extent of axon degeneration for individual unmyelinated axonal arbours over up to 3 days in the living adult mouse somatosensory cortex (**Fig. 1**). Our data provide new knowledge on cortical axon degeneration *in vivo:* Firstly, segments below ~ 600 μm, undergo a faster form of WD, supporting a protective factor level cut-off based model for cortical axon degeneration. Secondly, synaptic density but not arbor complexity, control cortical axon ROWD. Thirdly, AAD is negligible on cortical axons and does not affect the membrane in the spinal cord. Finally, similarly to WD in other fibre systems, a NAD^+^-dependent pathway independent of transcription regulates cortical axon ROWD.

### *In vivo* imaging assay to study single axon fibre degeneration in the mammalian brain

*In vivo* optical imaging of spinal cord injury has provided unique insights into the mechanisms of axon degeneration and regeneration in recent years (*35, 38, 72*). However, we are only now starting to exploit this tool to understand the principles of axon degeneration and regeneration in the mammalian brain (*48–50, 73–75*). Limitations of optical methods through glass windows to induce local damage include the inability to model more complex types of injury conditions such as those found in traumatic brain injury or multiple sclerosis and the difficulty of getting therapeutic compounds into the brain in order to manipulate the local environment of the neuropil *in vivo* (*76*). There are three main advantages of laser-mediated axonal transection compared to previously reported methods. Firstly, laser axotomy reveals the entire degeneration process as close as 4-10 μm from the injury site (*30, 49*). In contrast, due to anatomical distortions axonal changes can only be observed at a distance (>100 μm) from the injury site following mechanical or needlestick injury. Secondly, local tissue injury can be induced without rupturing the dura mater and overlying blood vessels, decreasing contributions from inflammatory mediators, thereby reducing the complexity of cellular interactions and improving experimental reproducibility (*49, 77*). Thirdly, several neurons, or even parts of a neuron (*54*), can be simultaneously targeted with minimal damage of the surrounding tissue (*30, 49, 51, 78–81*). The combination of this lesion method with the selective monitoring of the axolemma, rather than the axoplasm as in previous studies, allowed to accurately measure the timing of the initiation and execution phases of axon degeneration, ruling out instances of cytoplasmic fragmentation not directly accompanied by loss of membrane integrity (as we show it is the case for spinal cord AAD, **Suppl. Fig. 3**). We expect that this paradigm and its refinements, for example by using multi-colour imaging of cytoskeletal (*82*), mitochondria (*83*) and membrane markers, will be useful to investigate axonal degeneration in other mouse models of disease (e.g. diabetes and Alzheimer’s disease) or human-mouse chimeric models with patient-specific genetic background (*84*).

### Axon degeneration similarities and differences between fibre systems

We consider here the timing of axon fragmentation post-injury. This is the first study assessing the degeneration kinetics of individual axons in the mammalian brain. We used a labelling strategy targeted mainly to excitatory axons originating in cortical layer 2/3/5/6 and the thalamus, because of their vulnerability in several neurodegenerative and acute injury conditions. The reported kinetics fall within published time frames for WD in the murine spinal cord (*35*), peripheral nerves (*63*) and optic nerve (*2*), but reveal a surprising minor role for AAD in severed cortical axons (**Suppl. Fig. 2 and Table 1**). In addition, transected segments of unmyelinated cortical axons < ~ 600 μm undergo ROWD (**Fig. 2**), with different spatial and temporal progression patterns compared to both WD and AAD. Even though only a small percentage (< 1%) of lesioned axonal arbor length undergoes ROWD, ROWD affects up to 2-3 times more axon length than AAD (*35*) (**Table 1**).

For cortical axons the timing of both the initiation (i.e. onset, **Fig. 2d**, e) and execution phase (i.e. fragmentation kinetics, **Fig. 2d**, g) differ depending on a cut-off length of ~ 600 μm (**Table 1**). Our data rule out a simple linear length-based mechanism to explain the onset of cortical axon degeneration (*20*), and instead identify a cut off of ~ 600 μm, above which axons are protected for 1-2 days *in vivo*. The rapid onset for segments < ~ 600 μm can be explained by a cut-off based model, which postulates the existence of critical levels of a protective factor. In earlier work in zebrafish larvae, the length of the disconnected axonal segment was not a critical parameter, not affecting the duration of the lag phase and only slightly influencing the fragmentation rate (*30*).

The difference in fragmentation rates between WD and ROWD (**Fig. 2d**, g) could reflect the natural variability of the degeneration process across the nervous system. It is well established that degeneration kinetics are variable between axons in adults (*25*), and even more so during development. For example, axotomised trigeminal axons in the developing zebrafish show embryonic stage-dependent differences in the time of the degeneration onset and lack AAD (*30*). Other axon subtypes, such as those of the zebrafish lateral line projections, do undergo a process resembling AAD in that a gap rapidly forms on both sides of the lesion, although no evidence of fragmentation is detected (*39*).

### Mechanisms of cortical axon degeneration

Our results on the regulation of ROWD by NAD^+^ dependent mechanisms (**Fig. 4**) are consistent with previous data of delayed dopaminergic and striatal synaptic degeneration in Wld^S^ mice (*16, 85*), providing further *in vivo* evidence that Wld^S^-based neuroprotective therapeutics may be developed not only for myelinated but also for unmyelinated, grey matter fibre degeneration.

We find that axotomy-induced degeneration of damaged cortical axon terminals depends on synaptic density for segments < 600 μm undergoing ROWD, but not for segments undergoing WD (**Fig. 3**). Synapse loss in neurodegeneration can occur independently of axonal and cell body loss, suggesting that different mechanisms regulate the degeneration of these compartments (*46, 86, 87*). We hypothesise that for segments < 600 μm the presence of synapses limits the depletion of axon protective factors, prolonging the lag phase before fragmentation.

Unlike both WD and AAD of myelinated fibres where axoplasmic fragmentation commences either close to the lesion site or distally and then proceeds away from it (*25, 35*), axolemmal fragmentation of cortical axons did not always occur in an anterograde or retrograde direction (see Results section), challenging previous models where the fragmentation wave would be triggered by the clearance of putative protective factors via axonal transport (*20, 27*).

Together, our data provide evidence for the existence of a rapid-onset form of WD, which selectively affect segments < 600 μm (**Fig. 2**), is more likely to start from the distal end of the axon than WD and is influenced by the extent of synaptic density (**Fig. 3c**). We also uncovered a general mechanism of ROWD regulation involving a local, transcription-independent NAD^+^-mediated pathway in the damaged neurons (**Fig. 4**). In the future, the *in vivo* optical imaging approach used here could lead to a better understanding of the molecular and cellular mechanisms that lead to cortical axon ROWD, compared to WD or AAD, and could help to develop effective therapies to counteract or stabilise the progression of axon degeneration in both the diseased and injured brain.

## Acknowledgments

Massimo Hilliard, James Vickers, Shabana Khan, Michele Ettorre and Simone Di Giovanni provided comments to improve the manuscript. Arnau Hervera Abad for help with the laminectomy procedure. Raquel Real for help with the laser lesion experiments. Julia Hines for help with the processing of the time lapses. Thomas Carroll and Bill Bennett for help with statistical analysis. This work was supported by the Medical Research Council.

## METHODS

### Animals

Adult male mice from Thy1-L15 (*55*) (membrane-targeted GFP) (n = 16, 12-14 weeks) and the Wld^S^ line, which spontaneously expresses the slow Wallerian degeneration mutation (Wld^S^-Thy1-L15, *n* = 13, 12-14 weeks) were used for all *in vivo* imaging experiments unless otherwise specified. All mice were given access to food and water *ad libitum* and maintained in a 12 hour light-dark cycle. All procedures were conducted by researchers holding a UK personal license, conducted in accordance with the UK Animals (Scientific Procedures) Act 1986 and approved by the local Animal Welfare and Ethical Review Body (AWERB).

### Craniotomy

Cranial windows were surgically implanted over the somatosensory cortex as previously described (*88*). Mice were anaesthetized with ketamine-xylazine intraperitoneal injection (0.083 mg/g ketamine, 0.0078 mg/g xylazine) and then administered intramuscular dexamethasone (0.02 ml at 4 mg/ml), to reduce inflammation, and subcutaneous bupivacaine (1 mg/kg), a local anesthetic. A few drops of lidocaine (1 % solution) were applied on its skull surface prior to a 5 mm diameter craniotomy being drilled over the somatosensory cortex. A glass coverslip was then placed over the craniotomy and sealed with glue and dental cement. The skull was subsequently covered in dental cement and a metal bar placed on top for positioning at the two-photon microscope. Mice were allowed to recover for 14 days prior to imaging.

### Two-Photon imaging *in vivo*

A purpose built two-photon microscope (Prairie Technologies) equipped with a tunable coherent Ti:Sapphire laser (Coherent) and PrairieView acquisition software was used for all imaging experiments. Mice were anaesthetized with isofluorane (1-1.5 %) and secured to the microscope with the head metal bar attached to a custom-built fixed support. Lacri-lube was applied to the eyes to prevent dehydration and temperature maintained by a heating blanket. An Olympus 4X with a 0.13 numerical aperture (NA) objective was used to identify characteristic blood vessels to reliably locate regions-of-interest (ROIs) at each imaging time point. An Olympus 40X (NA = 0.8) water immersion objective was used to acquire several ROI stacks per animal (typically 100 x 100 μm field of view, 512 x 512 pixels, 1 μm step size). A pulsed 910 nm laser beam was used with typical power never exceeding 70mW on the back focal plane. Typically 1 - 4 axons were chosen per animal. Each imaging session lasted a maximum of 90 minutes, after which animals recovered for at least 30 minutes before the next session began.

### Laser mediated axonal lesions *in vivo*

A total of 122 axons (58 *in vivo* and 64 *in vitro*) were lesioned and followed for up to 72h *in vivo* and 96 h *in vitro* to assess degeneration kinetics. Axons located in the upper layers of the cortex were selected starting from their tips/endings, then followed along the shaft towards the cell body for as long as axon identification could be ascertained, until the axon disappeared under a blood vessel, under the bone edges or could not be distinguished from neighbouring axons, which defined axonal segments of variable length (**Fig. 1**-2). A laser micro-lesion protocol was optimized to induce a reproducible lesion as in (*49, 50*). Briefly, the imaging laser was tuned to 800 nm and engineered to deliver a single spot scan to a previously identified axon, severing the axon shaft. Lesions were conducted in layer-1 of the cortex within 50 μm of the pial surface. Care was taken to lesion away from large vessels to avoid any confounding effects of the breakdown of the blood-brain-barrier. Complete axotomy was confirmed in each case by the eventual disappearance of the disconnected distal axon segment.

Since it was not possible to predict when the axon fragmentation process would start, cortical axons were imaged before the lesion and at variable intervals after the lesion, until at least 50% of the disconnected axon had disappeared. 89.6 ± 13.9% of axon length disappeared over 5.5 time points per axon (range 3-21 imaging sessions, *n* = 43 *in vivo* cortical axons). To ensure the welfare of the animals over the prolonged imaging sessions, which lasted up to 3 days (**Fig. 2d**), while still ensuring accurate measurements of the fragmentation kinetics, imaging intervals were kept to less than four hours between sessions (i.e. the maximum interval between any two consecutive sessions was 4 hours). As a result, the average time interval recorded after fragmentation onset was 103 min, with a minimum of 3 images collected during the fragmentation phase for any axon (average 4 images, range 3-10). Therefore, the maximum fragmentation speed we can measure on average for a segment undergoing ROWD is ~ 300 μm / 103 min = ~ 3 μm/min, and for a segment undergoing WD, ~ 1000 / 103 = ~ 10 μm/min, i.e. when the axonal segment is entirely removed between two consecutive time points. Reducing the average imaging interval after fragmentation onset by a factor of ~ 10, to 13 min (with a minimum of 4 images collected during the fragmentation phase for any axon; average 9 images, range 4-16), however, did not change significantly the measured ROWD fragmentation rates (0.6 ± 0.1 μm/min; *n* = 8; *P* > 0.05 compared to segments < 600 μm imaged on average every 103 min) and onset (160 ± 24 min; *n* = 8 segments imaged with less than 2 hour intervals; *P* > 0.05 compared to segments < 600 μm imaged with less than 4 hour intervals), confirming our methodology. A subset of axons (*n* = 16 imaged with a membrane marker and 10 with an axoplasm one) was imaged over minutes for the first 60 minutes to investigate AAD (**Supplementary Fig. 2**). These experiments, which consisted of higher number of imaging time points acquired, also rule out any contribution of photo-bleaching of membrane-bound GFP to the interpretation of degeneration kinetics. After the imaging, mice were administered a lethal dose of ketamine/xylazine and transcardially perfused with ice-cold saline and followed by paraformaldehyde. Brains were removed and post-fixed overnight in paraformaldehyde at 4°C before being transferred to 0.01 M PBS and stored in the fridge.

### Laminectomy and spinal cord imaging

Dorsal laminectomy was performed at spinal levels L1-L3. Mice (n = 10) were anaesthetized with intraperitoneal injection of ketamine-xylazine mixture (0.083mg/g ketamine, 0.0078 mg/g xylazine). After shaving the back, a longitudinal incision was made to retract the skin. The perivertebral muscles were deflected to expose the vertebral column and the laminae were removed using curved blade scissors. The mice were then moved carefully to the microscope stage. The trunk was raised slightly and the vertebral column was clamped between two custom made bars for stabilization during 2-photon imaging. The area surrounding the exposed spinal cord was circled by a layer of veterinary n-butyl cyanoacrylate adhesive to facilitate the formation of a saline meniscus between the objective and the specimen. Labelled ascending axons in the dorsal white matter tracts were targeted for micro-lesion. The anesthesia level was maintained by inhalation of an oxygen and isoflurane mixture (1.5 - 2.5%) throughout the imaging sessions.

### Organotypic cortical slice culture preparation

Organotypic slices were prepared from the brains of neonatal transgenic mice (Thy1 - L15, described above) based on (*69*). Pups (n = 29) were sacrificed at postnatal day (P) 3-4 by decapitation. Brains were carefully removed and immediately submerged in ice-cold culture medium (MEM Hanks supplemented with 20% heat-inactivated horse serum, Penicillin/Streptomycin (1 %), Insulin (1 μg/ml), CaCL2 (1 mM), MgSO4 (2mM), ascorbic acid (1.2 %)). The cortical hemispheres were dissected away from the diencephalon and cortical slices (400 μm) were obtained with a McIlwain Tissue Chopper (Laboratory Engineering Co. Ltd, UK). Slices were carefully transferred, using a fire-polished, wide bore glass Pasteur pipette, to 35 mm petri dishes containing fresh, ice-cold culture medium and incubated on ice for 40 minutes. Using a dissection microscope, slices containing the somatosensory cortex were carefully transferred to Millicell culture plate inserts (Millipore) and incubated for up to 2 months with 1 ml culture medium at 35°C in a humidified atmosphere with 5% CO2. The culture medium was changed twice weekly throughout the culture period.

### Imaging of axon degeneration in organotypic slice cultures

Organotypic slice cultures were imaged using two-photon microscopy beginning at 21-28 days *in vitro* (DIV). Culture inserts containing slices were transferred to sterile 35 mm petri dishes containing 1 ml pre-warmed Tyrodes salt solution (Sigma) and a further 1 ml was added on top of the slice. Imaging sessions of up to 1.5 hours, using low power (< 20 mW at the back focal plane) were possible without noticeable deleterious effects in the slice. Typically 1-4 axons were chosen per slice (maximum length, 5 mm). Laser-mediated micro-lesions were performed as described for *in vivo* imaging protocols. To investigate the effects of NAD^+^ on the dynamics of axonal degeneration, slices were incubated for 24 hours pre- or postlesion, with medium containing 1 mM NAD^+^ or with medium alone. Following imaging and/or lesioning, the salt solution was carefully removed from above and below the membrane and the inserts were returned to the incubator with fresh, prewarmed culture medium (with or without NAD^+^, as appropriate).

### Data Analysis

Maximum projections of individual image stacks from *in vivo* and *in vitro* two-photon images of axons in WT and Wld^S^ mice were processed with ImageJ (file conversions, length quantifications) and Adobe Photoshop (section alignment, image rotation, brightness and contrast adjustment).

Synapse density was determined by annotating *Terminaux* Boutons (TBs) and *En Passant* Boutons (EPBs) in Matlab using custom made software (*89, 90*. TBs (defined as protrusions between 1–5 microns) and EPBs (defined as swellings with pixel intensity two times brighter than the axon backbone) were scored according to stringent criteria based on (*88*) and (*89*).

The length of the distal portions from lesion to ending was measured for all axons. ‘Distal’ and ‘close to the lesion’ ends were defined as those within 100 μm from the ending or lesion site respectively (**Fig. 1d**). Axons were split into unbranched and branched axons where the total length was less or more than 600 μm. To determine the onset and rate of fragmentation, severed axons were imaged at regular intervals of less than four hours (or for a subset of 8 axons less than 2 hours, see above) and the time point of the imaging session immediately preceding the first sign of clearly visible break in the axon (i.e. fragmentation) recorded as the onset time. A decrease in length of more than 10 μm over two consecutive sessions was considered as degeneration. At each subsequent imaging session, the sum of the individual fragment lengths was expressed as a percentage of the total unlesioned length of the axon. The gradient (m) of the resulting slope (**Fig. 1g**) was used to calculate the fragmentation rate using the formula y = mx + c (where c is the y intercept).

Figures were prepared using Microsoft Office Suite and Adobe Illustrator. Statistical analyses were performed in Microsoft Excel, R or GraphPad Prism. Unless otherwise stated, all measurements are given as the mean ± standard error of the mean (SEM). Student’s t-tests or Mann Whitney U tests were performed, unless otherwise stated. Results were considered significant when *P* < 0.05.

---

**Supplementary Fig. 1.**
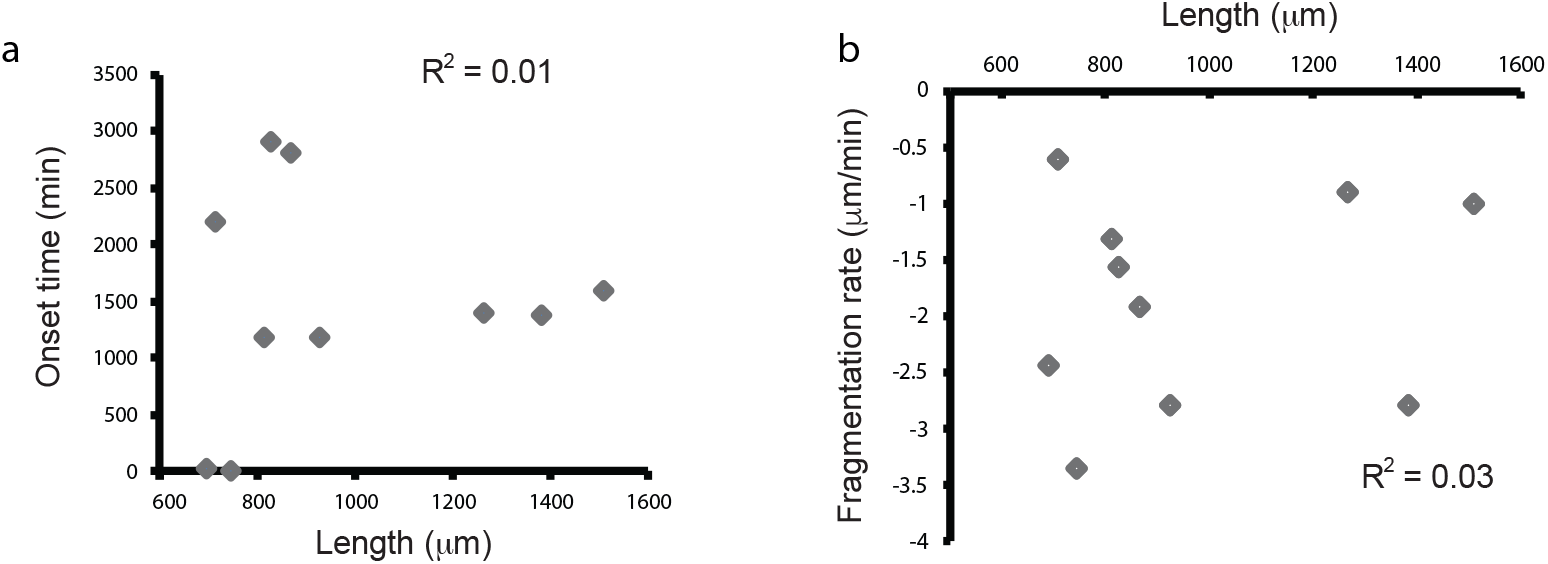
No correlation between axon length and WD kinetics. (**a**) No correlation between onset time and length for 10 axons undergoing WD (P > 0.05). (**b**) No correlation between fragmentation rate and length for 10 axons undergoing WD (*P* > 0.05).

**Supplementary Fig. 2.**
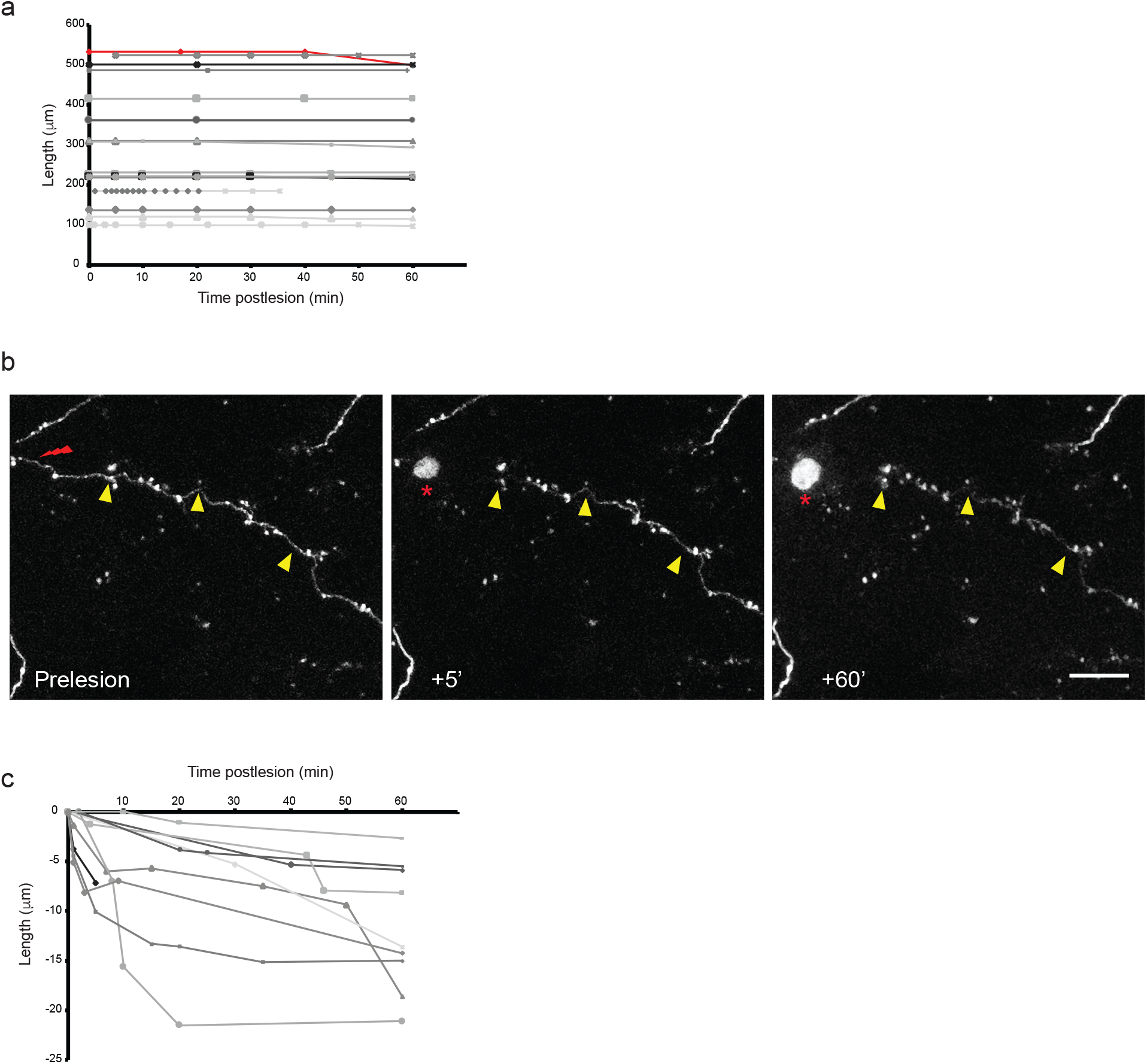
AAD is negligible in injured cortical axons *in vivo*. (**a**) Length of severed cortical axon distal segments monitored with a membrane marker (L15 (*55*)) over time *in vivo*. Note length stability in all but one axon (red line), indicating absence of fragmentation over the first 60 minutes postlesion (*n* = 16 axons). (**b**) Example of 2-photon *in vivo* imaging of distal axon apparent retraction from the lesion site monitored with cytosolic markers (GFP-M line (*60*)). Note the absence of fragmentation over the first 60 minutes postlesion. Scale bar = 10 μm. (**c**) Quantification of the distance from the lesion site to the distal axon stump at the indicated postlesion times. *n* = 10 axons monitored with a cytosolic marker.

**Supplementary Fig. 3.**
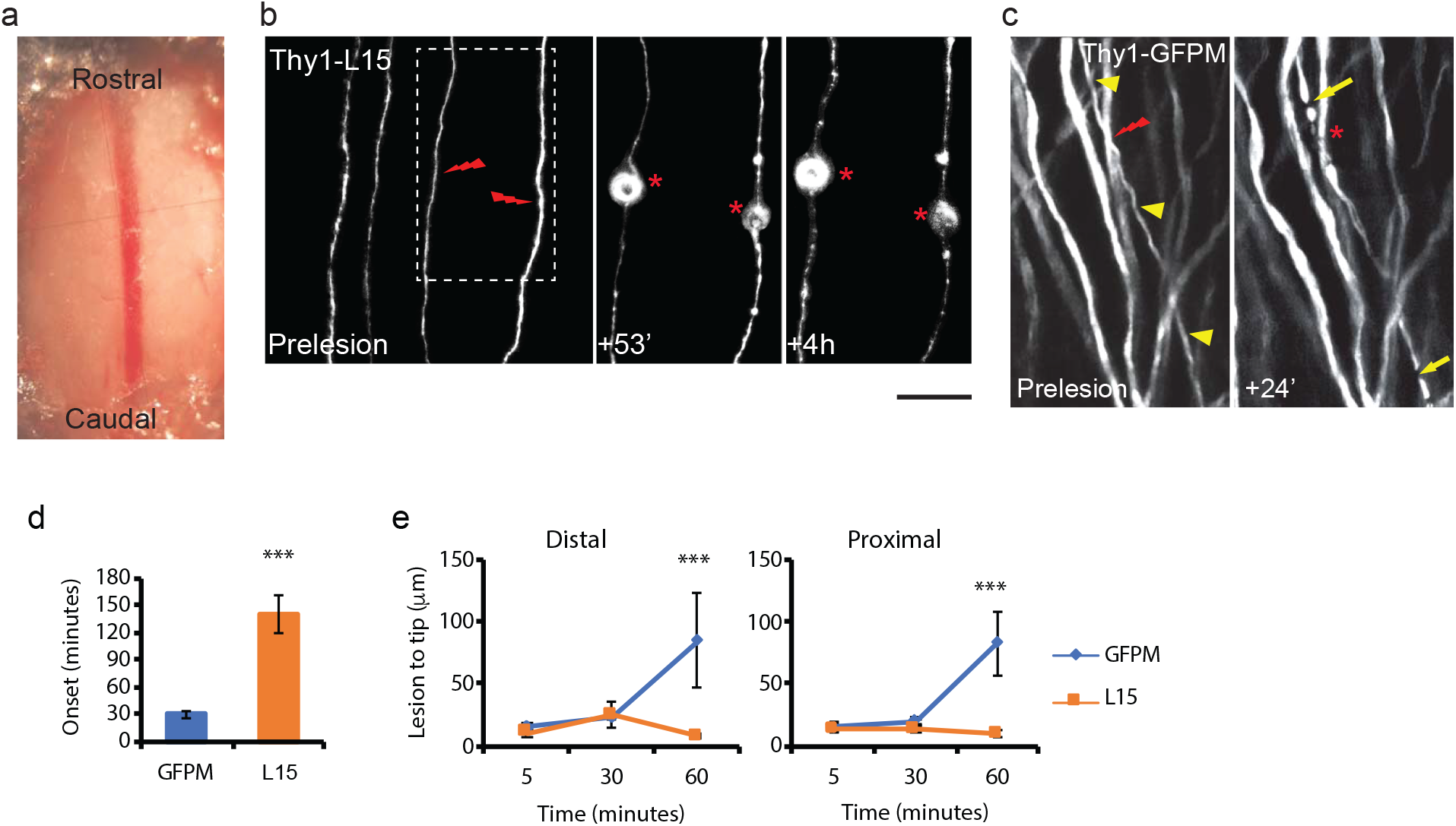
AAD involves axoplasmic but not axolemmal fragmentation in spinal cord sensory axons after axotomy. (**a**) Representative segment of dorsal spinal cord exposed after laminectomy. (**b**) No fragmentation up to 4 h post lesion in Thy1 - L15 sensory axons expressing a membrane-bound form of GFP (n = 9 axons, 5 mice). Note beading, but no fragments up to 4 hours postlesion. Lightning bolt and asterisks indicate lesion site. The area highlighted by the white dashed box is magnified in the postlesion panels. (**c**) AAD (arrows) after laser-mediated lesions observed in sensory neurons expressing a cytosolic form of GFP (n = 9 axons, 5 mice). Lesioned axon indicated by arrowheads. Lightning bolt and asterisks indicate lesion site; arrows indicate axoplasmic fragmentation in Thy1-GFP-M, 24 min after axotomy. (**d**) The mean onset of sensory axon fragmentation is significantly shorter for GFP-M (cytosolic GFP), compared to L15 (membrane-targeted GFP) axons. (**e**) Absence of membrane fragmentation in AAD up to 1 hour postlesion. Scale bar is 1 mm in (a), 50 μm in (b) and in (c).

**Supplementary Movie 1.** Time-lapse movie of axon degeneration in the living mouse brain.

